# Quantitative Cross-Species Comparison of Serum Albumin Binding of Per- and Polyfluoroalkyl Substances from Five Structural Classes

**DOI:** 10.1101/2023.11.10.566613

**Authors:** Hannah M. Starnes, Thomas W. Jackson, Kylie D. Rock, Scott M. Belcher

**Affiliations:** Department of Biological Sciences, North Carolina State University, 127 David Clark Labs Campus Box 7617, Raleigh, NC 27607, USA; Public Health and Integrated Toxicology Division, Center for Public Health and Environmental Assessment, U.S. Environmental Protection Agency, Research Triangle Park, NC 27711, USA; Department of Biological Sciences, Clemson University, Clemson, SC 29634, USA

**Keywords:** binding affinity, differential scanning fluorimetry, *in vitro*, PFAS, species extrapolation

## Abstract

Per- and polyfluoroalkyl substances (PFAS) are a class of over 8,000 chemicals that are persistent, bioaccumulative, and toxic to humans, livestock, and wildlife. Serum protein binding affinity is instrumental in understanding PFAS toxicity, yet experimental binding data is limited to only a few PFAS congeners. Previously, we demonstrated the usefulness of a high-throughput, *in vitro* differential scanning fluorimetry assay for determination of relative binding affinities of human serum albumin for 24 PFAS congeners from 6 chemical classes. In the current study, we used this differential scanning fluorimetry assay to comparatively examine differences in human, bovine, porcine, and rat serum albumin binding of 8 structurally informative PFAS congeners from 5 chemical classes. With the exception of the fluorotelomer alcohol 1H,1H,2H,2H-perfluorooctanol (6:2 FTOH), each PFAS congener bound by human serum albumin was also bound by bovine, porcine, and rat serum albumin. The critical role of the charged functional headgroup in albumin binding was supported by the inability of serum albumin of each species tested to bind 6:2 FTOH. Significant interspecies differences in serum albumin binding affinities were identified for each of the bound PFAS congeners. Relative to human albumin, perfluoroalkyl carboxylic and sulfonic acids were bound with greater affinity by porcine and rat serum albumin, and perfluoroalkyl ether congeners bound with lower affinity to porcine and bovine serum albumin. These comparative affinity data for PFAS binding by serum albumin from human, experimental model and livestock species reduce critical interspecies uncertainty and improve accuracy of predictive toxicity assessments for PFAS.

## Introduction

Per- and polyfluoroalkyl substances (PFAS) are a class of synthetic chemicals, comprising thousands of structurally diverse compounds with at least one fully fluorinated methyl or methylene group (OECD 2021). Due to their stability, persistence, and widespread use, PFAS are ubiquitously detected in biological and environmental matrices (Cousins et al. 2020; Guillette et al. 2020; Kotlarz et al. 2020; Cousins et al. 2022; Guillette et al. 2022; Kirkwood et al. 2022; Rock et al. 2023). Some PFAS are toxic to multiple organ systems, including the hepatic, renal, immune, reproductive and nervous systems (Carlson et al. 2022; Radke et al. 2022; Crawford et al. 2023). Within the body many PFAS partition to, and accumulate in, protein-rich tissues, with highest concentrations found in the blood, liver, and kidneys of most exposed species (Hoff et al. 2003; Bischel et al. 2011). The majority of PFAS research has focused on perfluorooctanesulfonic acid (PFOS) and perfluorooctanoic acid (PFOA), whereas physiochemical properties of thousands of chemicals in this heterogeneous class remain uncharacterized (Radke et al. 2022).

Serum albumin is the most abundant protein in vertebrate blood, constituting over 50% of human blood proteins (Peters 1995; Merlot et al. 2014; Raoufinia et al. 2016). Serum albumin is the primary blood transport protein for many endogenous and exogenous ligands, including but not limited to long-chain fatty acids, bile acids, bilirubin, steroids, hormones, ions, and many pharmaceuticals (Peters 1995; Merlot et al. 2014). Additionally, several perfluoroalkyl acids (PFAAs), including PFOS, PFOA, perfluorohexanesulfonic acid (PFHxS), perflurononanoic acid (PFNA), and perfluorodecanoic acid (PFDA), are bound and transported by albumin in human blood (Forsthuber et al. 2020). Experimental toxicokinetic studies in animals have demonstrated that PFOS, PFOA, and PFHxS are similarly bound by serum albumin of cynomolgus monkeys and rats, at affinities comparable to those of many endogenous ligands and pharmaceuticals (Kerstner-Wood et al. 2003; Varshney et al. 2010; U. S. EPA 2016a; U. S. EPA 2016b).

In addition to influencing ligand transport and distribution, albumin binding affinity is an important determinant of biological half-life and bioavailability for many ligands (Kragh-Hansen 1981; Peters 1995; Sleep et al. 2013; Lexa et al. 2014; Merlot et al. 2014; Zorzi et al. 2019). Increased albumin binding affinity has a pivotal role in prolonging the half-life of many bioactive compounds (Sleep et al. 2013). Along with increasing biological half-life and delaying elimination, plasma protein binding also decreases the relative free-fraction of PFAS available for uptake in target tissues, highlighting the potentially multifaceted role of albumin-PFAS interactions in determining pharmacokinetic properties of individual PFAS congeners (Kragh-Hansen 1981; Peters 1995; Han et al. 2003; Ascenzi and Fasano 2010; Merlot et al. 2014; Sheng et al. 2020).

Physiologically based pharmacokinetic (PBPK) modeling conducted by Cheng & Ng (2017) has suggested that albumin concentration and binding affinity are among the most important predictive parameters in determining overall PFOA blood concentration and tissue distribution in the rat. Additional PBPK modeling studies found that the plasma free fraction of PFOS and PFOA is most likely to travel between body compartments, and is therefore a key determinant of cellular uptake (Fàbrega et al. 2014; Cheng and Ng 2017; Chou and Lin 2019; Chou and Lin 2021; Deepika et al. 2021). For example, a study conducted by Gao et al. (2019) found an inverse correlation between binding affinity of HSA for PFAS congeners and placental transfer efficiency.

Although advanced computational approaches have been developed to predict PFAS bioactivity and toxicity, there is significant uncertainty in the results of these predictive models due to the lack of foundational physiochemical data describing PFAS-protein interactions (Cheng and Ng 2017; Cheng and Ng 2018; Cheng and Ng 2019). To improve quantitative binding predictions, physiologically-based toxicokinetic models, and ultimately risk assessments that inform exposure limits to protect public health, there is a critical need for experimental data describing albumin-PFAS interactions with structurally diverse sets of PFAS (Fenton et al. 2021).

In addition to a lack of physiochemical data, there are also translational data gaps that limit ability to extrapolate PFAS toxicity data from animal models to humans. Major cross-species differences exist in PFAS elimination rates, with half-lives for some PFAS that are orders of magnitude apart in humans compared to model organisms and livestock (Pizzurro et al. 2019; Fenton et al. 2021). While estimates vary across studies, reported half-lives for PFOS and PFOA tend to be on the order of hours to weeks in rodents, days to months in livestock, and years in humans (Russell et al. 2013; Numata et al. 2014; Lau 2015; Pizzurro et al. 2019; Death et al. 2021). As albumin binding plays a key role in regulating PFAS bioaccumulation and distribution, differences in albumin-PFAS binding may influence differences in biological half-lives across species (D’eon et al. 2010; Cheng et al. 2021; Fenton et al. 2021).

Differential scanning fluorimetry (DSF) is an *in vitro* method used to compare changes in thermal denaturation of ligand-free and ligand-bound proteins that accurately defines binding affinities for protein-PFAS interactions (Niesen et al. 2007; Jackson et al. 2021). Along with being fit-for purpose, the DSF assay is scalable to high-throughput formats, requires small reagent volumes, and is cost-effective, a combination of properties that are advantageous compared to other methods for measuring protein-PFAS binding affinity, including isothermal titration calorimetry, surface plasmon resonance, or mass spectrometry (Niesen et al. 2007; Vivoli et al. 2014; Liu et al. 2019; Jackson et al. 2021).

We previously demonstrated the usefulness of DSF for evaluating human serum albumin (HSA) binding of PFAS, reporting relative binding affinities for 24 PFAS from 6 chemical classes (Jackson et al. 2021). Using this DSF assay, we identified differences in binding mediated by differences in PFAS functional groups and subtle changes in perfluoroalkyl chain length, demonstrating the capability and sensitivity of this assay for determining binding affinity differences among closely related congeners (Jackson et al. 2021). Due to the combined benefits of assay sensitivity and scalability for high-throughput analyses, this DSF assay is capable of filling critical gaps in describing the physiochemical properties of PFAS (Jackson et al. 2021).

To address the lack of comparative species specific PFAS-protein binding data, we utilized DSF to comparatively evaluate relative binding affinities of serum albumin from human, laboratory rat, and livestock (cow and pig) species for a subset of 8 structurally informative PFAS. The PFAS analyzed included the four congeners with completed toxicity assessments by the US Environmental Protection Agency (EPA): PFOA, PFOS, perfluorobutanesulfonic acid (PFBS) and hexafluoropropylene oxide-dimer acid (HFPO-DA) (U.S. EPA, 2016, 2021b, 2021a). To evaluate the generalizability of trends observed in HSA-PFAS binding based on carbon chain length, perfluorocarboxylic acid (PFCA) and perfluorosulfonic acid (PFSA) congeners with 4- and 8-carbon alkyl chains were included. Further, we leveraged two computational tools, the EPA Sequence Alignment to Predict Across Species Susceptibility (SeqAPASS) application and a molecular docking approach, to identify species-specific amino acid sequence differences in serum albumin and to gain predictive insights into the mechanistic basis for observed interspecies differences in albumin-PFAS binding affinities (LaLone et al. 2016; Cheng et al. 2021).

## Materials and Methods

### Chemicals and Reagents

Aqueous solutions were prepared with sterile Milli-Q A10 water (18 Ω; 3 ppb total oxidizable organics). GloMelt (λEx = 468, λEm = 507 nm) and carboxyrhodamine (ROX; λEx = 588, λEm = 608 nm) dyes (CAT #33022-1) were purchased from Biotium (Fremont, CA). Bovine serum albumin (BSA, purity ≥ 98%, fraction V, BP1605, Lot 053997), methanol (purity ≥ 99.9%, CAT #A454-4), dimethyl sulfoxide (purity ≥ 99.9%, CAT #136-1, Lot 147002), Na_2_HPO_4_ (purity ≥ 99.2%, CAT #S374-3, Lot 056560), NaCl (purity ≥ 100%, CAT #S271-10, Lot 134874), and 4-(2-(2-hydroxyethyl)-1-piperazineethanesulfonic acid (HEPES, purity ≥ 99.%, CAT #BP310-1, Lot 052975) were from Thermo Fisher Scientific (Waltham, MA). Rat serum albumin (RSA, purity ≥ 99%, A6414, Lot SLCF8334), and porcine serum albumin (PSA, purity ≥ 98%, A1830, Lot SLBZ2093) were from Sigma-Aldrich (St Louis, MO). Octanoic acid (OA, CAS 124-07-2, purity ≥ 95%) was from Alfa Aesar (Havermill, MA). Structures and chemical classes of the PFAS congeners analyzed are shown in Figure 1.

**Figure 1.**
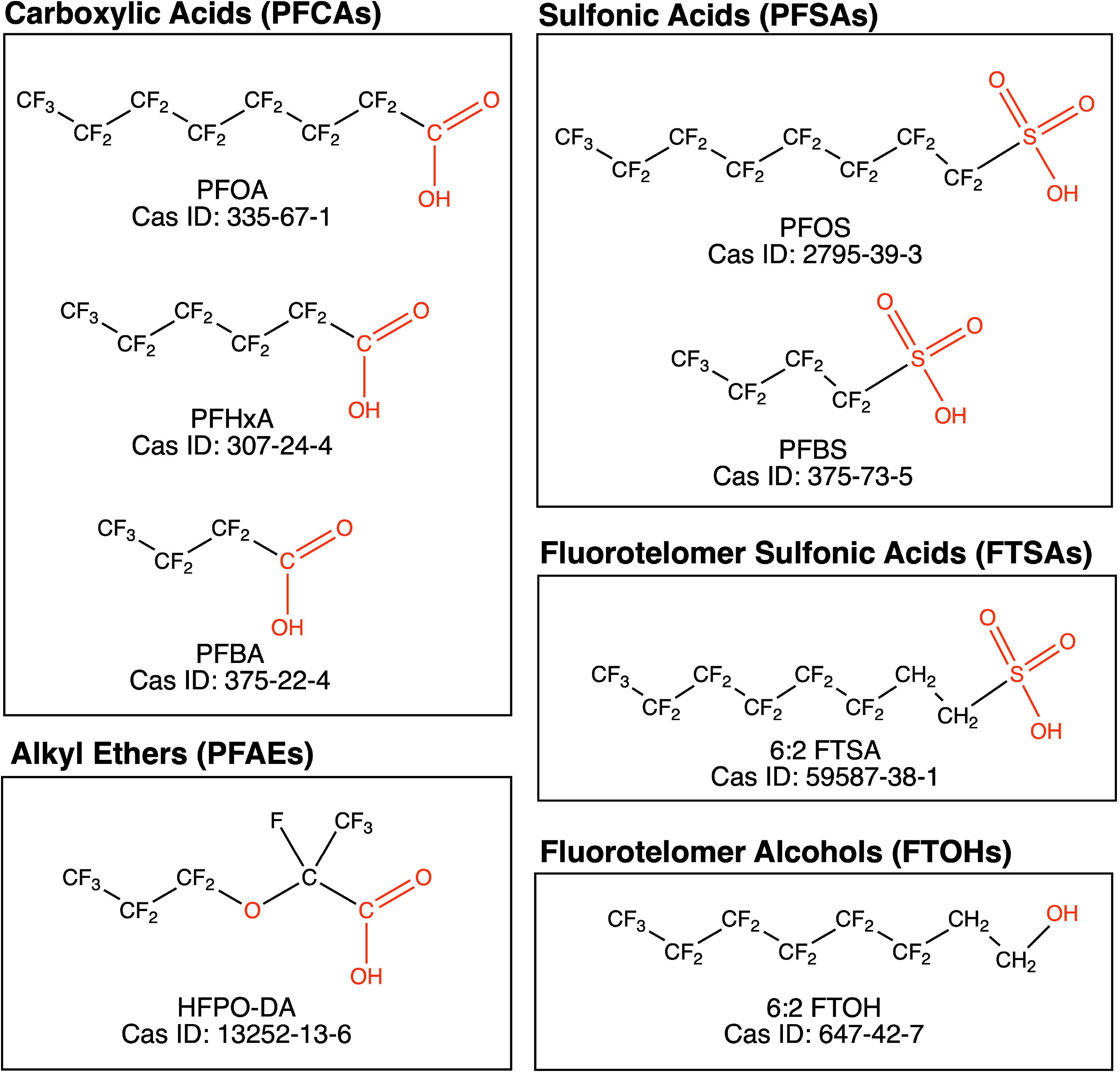
Chemical structures and Chemical Abstracts Service (CAS) registration numbers of PFAS congeners analyzed.

Perfluorobutanoic acid (PFBA, CAS 375-22-4, purity ≥ 99%), PFOA (CAS 335-67-1, purity ≥ 95%), and HFPO-DA/GenX (CAS 1352-13-6, purity ≥ 97%) were from Alfa Aesar (Havermill, MA). Perfluorohexanoic acid (PFHxA, CAS 307-24-2, purity ≥ 98%) and PFBS (CAS 375-73-5, purity ≥ 98%) were from TCI America (Portland, OR). PFOS (CAS 2795-39-3, purity ≥ 98%) was from Matrix Scientific (Columbia, SC), and 1H,1H,2H,2H-perfluorooctanol (6:2 FTOH, CAS 647-42-7, purity ≥ 97%) and 2H,2H,3H,3H-perfluorooctane-1-sulfonate (6:2 FTSA, CAS 59587-39-2, purity ≥ 97%) were from Synquest Laboratories (Alachua, FL).

### PFAS and Albumin Preparation

PFAS stock solutions (10 or 20 mM) were prepared in aqueous HEPES-buffered saline (HBS; final concentrations were 140 mM NaCl, 50 mM HEPES, 0.38 mM Na_2_HPO_4_, pH 7.4) and appropriate solvent when necessary (Table 1). Serum albumin stocks (1 mM) were prepared by reconstituting lyophilized stocks in HBS, with expected protein concentrations confirmed using the colorimetric Bio-Rad DC Protein Assay (Hercules, CA).

**Table 1.**
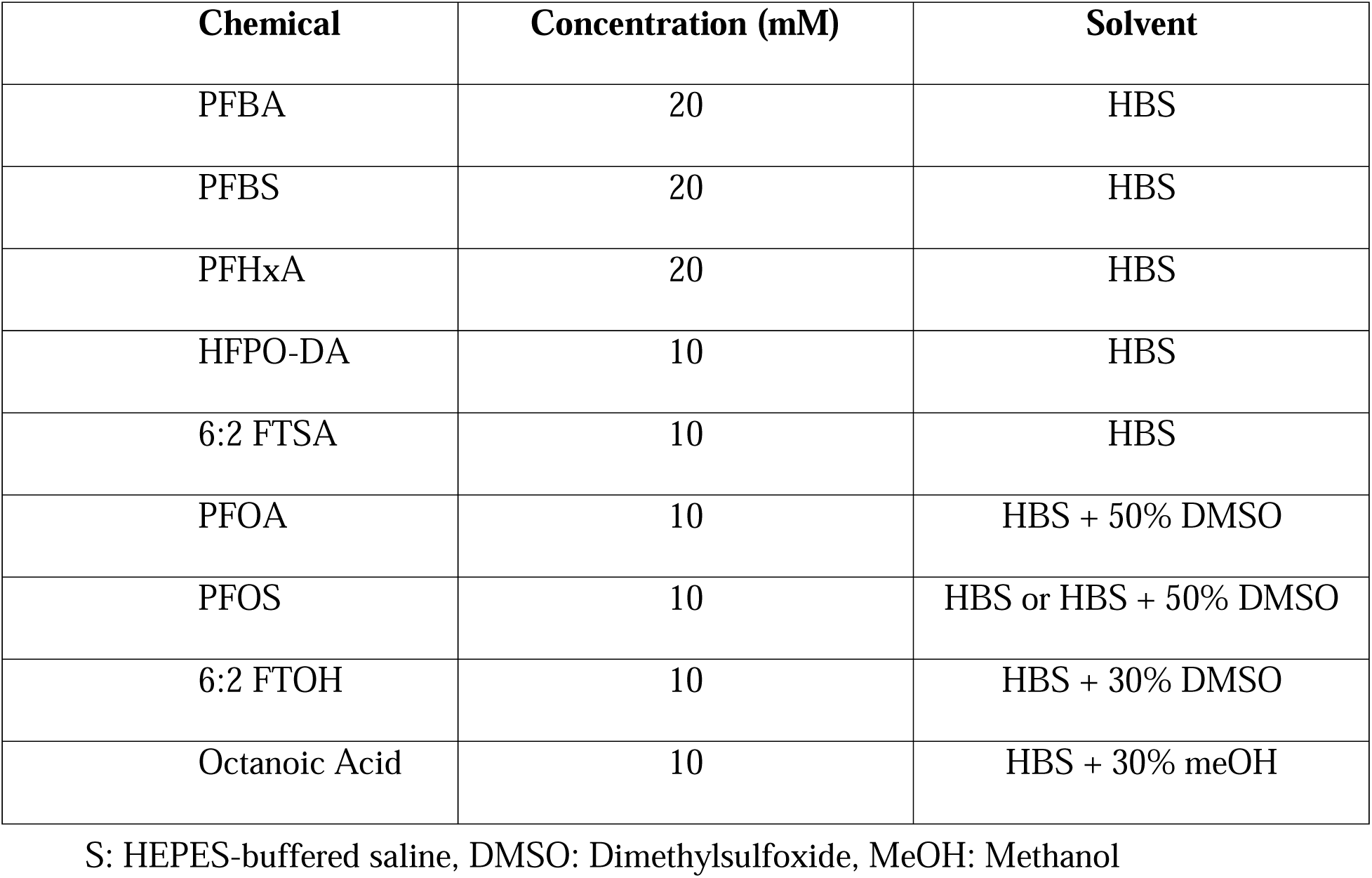
Test Chemical Stocks HB.

### Differential Scanning Fluorimetry Thermal Shift Assays

Individual assays were performed in sealed optical 96-well reaction plates (MicroAmp Fast, Applied Biosystems) at a final volume of 20 μL as previously described (Jackson et al., 2021). A minimum of two independent plates were run for each protein-ligand combination, with three replicates per sample on each plate. Controls on each plate included a no protein control, matching vehicle control (no ligand), and three concentrations of OA (1, 2, 3 mM) and/or PFHxA (0.3, 1, 3 mM) as positive controls. The concentration of albumin that yielded maximal signal-to-noise ratio was optimized for each species. Final protein concentrations for each assay were 0.13 mM for HSA, 0.18 mM for BSA, 0.11 mM for PSA, and 0.18 mM for RSA. For concentration-response analyses, PFAS stock solutions were serially diluted into HBS. **Sequence Alignment to Predict Across Species Susceptibility**

The SeqAPASS tool (https://seqapass.epa.gov/seqapass/; v. 6.1) was used to compare albumin primary amino acid sequences across species, with more targeted analysis comparing conservation of specific amino acid residues involved in PFAS binding (LaLone et al. 2016). For level 1 analysis, HSA (NCBI AAA98797.1) was used as the query protein to assess conservation of the complete amino acid sequence of BSA (NCBI P02769.4), PSA (NCBI ABM92961.1), and RSA (NCBI NP_599153.2). Level 3 analysis evaluated cross-species conservation of key amino acid residues in Sudlow sites I and II (Sudlow et al. 1975; Sudlow et al. 1976; Chen and Guo 2009; Ascenzi and Fasano 2010). An amino acid substitution was identified in the Sudlow I site of BSA compared to HSA, and was further investigated using the DUET web-based tool (https://biosig.lab.uq.edu.au/duet/stability), which predicts the impacts of specified amino acid mutations on protein stability (Pires et al. 2014; Cheng et al. 2021). The crystal structure for HSA (PDB ID 1AO6) was submitted to DUET, and a mutation was specified to alter the amino acid residue at site 211 from phenylalanine (found in HSA) to leucine (found in BSA) in Sudlow site I of BSA.

### Molecular Docking

To further investigate interspecies differences in albumin-PFAS binding, Autodock Vina (v. 1.2.0) was used for molecular docking analyses (Trott and Olson 2009; Eberhardt et al. 2021). The RCSB Protein Data Bank (PDB; https://www.rcsb.org/) was used to obtain high resolution (≤ 2.5 Å) three-dimensional protein structures. To assess HSA-PFAS docking, we used three monomeric structures previously chosen by Ng and Hungerbühler (2015) to evaluate HSA-PFAA docking: HSA complexed with myristic acid (1E7G), HSA complexed with PFOS (4E99), and an intermediate conformation between the two (1H9Z) (Bhattacharya et al. 2000; Petitpas et al. 2001; Luo et al. 2012; Ng and Hungerbuehler 2015). Only one high-resolution BSA structure was available in PDB (4F5S), which was selected for use in our experiments (Bujacz 2012). As 4F5S is a dimeric structure, one of the monomers was removed from the PDB file before docking. As of the date of publication, there were no crystal structures of PSA or RSA available in PDB. For comparative purposes, we followed the protocol for molecular docking described by Ng and Hungerbühler (2015), with the exception that PFAS ligand files were downloaded as three-dimensional SDF conformer files from NCBI PubChem database (https://pubchem.ncbi.nlm.nih.gov/), and converted to PDB format using PyMOL (Schrödinger L.L.C. 2017). Protein and ligand structures were prepared for docking using AutoDock Tools (v.1.5.7) as previously described (Morris et al. 2009; Trott and Olson 2009). Binding site boundary dimensions for six sites, defined by Ng and Hungerbühler (2015), were selected using the Grid Box feature in Autodock Tools.

To evaluate experimental success of docking, we redocked PFOS to the crystallized HSA protein (PDB 4E99), and used PyMOL to measure the Root Mean Square Deviation (RMSD) between our docking prediction and the ligand position in the crystal structure (Ng and Hungerbuehler 2015; Schrödinger L.L.C. 2017; Cheng and Ng 2018). Calculated RMSD values below 2 Å confirmed that Autodock Vina was able to successfully predict binding conformations for HSA and PFOS (Ng and Hungerbuehler 2015; Li et al. 2021).

### Data Analysis and Statistics

Raw DSF assay data, reported in relative fluorescent light units (RFU), were exported to Excel (Microsoft), and statistical analyses were performed using Prism (v.9.4.0, GraphPad Software Inc., San Diego, CA), SPSS v28 (IBM, Armonk, NY), or R statistical environment (R Core Team 2022). Melting temperature (T_M_) of albumin was defined as the temperature at which the maximum change in fluorescence occurs (Jackson et al. 2021). PFAS concentration-response curves were smoothed (Savitzky and Golay 1964), and dissociation constant (K_d_) values calculated using a single-site ligand binding model previously described (Vivoli et al. 2014; Jackson et al. 2021). For the biphasic shift in melting temperature observed for BSA binding of PFOS, the percent occupancy was used to calculate binding affinity (Vivoli et al. 2014).

A Kruskal-Wallis test with Dunn’s multiple comparisons test was used to compare the baseline melting temperatures of albumins. Generalized linear modeling (GLM) with a log-link Gaussian distribution was used for comparative evaluation of binding affinities, with PFAS and species as fixed factors, in the *lme4* R package (Bates et al. 2015). McFadden’s pseudo-R^2^ was calculated to evaluate goodness of fit for the GLM (Veall and Zimmermann 1996). Effect sizes for main effects of PFAS, species, and PFAS x species interaction on binding affinity were calculated using the *domin* function of the *domir* statistical package in R, reporting a general dominance statistic (GD) that represents the proportion of variance associated with each main effect in the model (Luchman 2023). The *emmeans* R package was used to evaluate contrasts of interest, with Sidak’s multiple comparisons correction, to compare K_d_ differences between HSA and other albumins for each PFAS congener, and to evaluate the intraspecies effect of carbon chain length on K_d_ by comparing binding of PFBA and PFOA, and PFBS and PFOS (Lenth 2023). For tests in which differences between binding constants were found, effect sizes for the post-hoc analyses were determined by calculating Cohen’s d (Lakens 2013). Effect sizes described by Cohen’s d are defined as small if d = .2, medium if d = .5, and large if d = .8 (Lakens 2013). For all statistical analyses, significant differences in means are defined as p < .05.

For each protein-PFAS combination, molecular docking ΔG predictions for the top 9 binding modes in each of the six analyzed binding sites were averaged. The geometric mean was taken across all binding sites for each protein, to represent the average ΔG of binding for that protein to each PFAS congener. To assess the strength of the linear relationship between calculated K_d_ values and molecular docking ΔG predictions for each protein-PFAS combination, Pearson’s correlation coefficients were computed using GraphPad Prism.

## Results

### Species Comparison of Serum Albumin Melting Temperatures

In the absence of ligand, the observed average T_M_ ± SEM for HSA was 71.3 ± 0.22°C, BSA was 62.6 ± 0.04°C, PSA was 71.3 ± 0.09°C, and RSA was 66.8 ± 0.18°C. A Kruskal-Wallis test with Dunn’s multiple comparisons test (H = 184.8, p < .0001) revealed that baseline melting temperatures across each albumin protein were significantly different from one another, with the exception of HSA and PSA (p > .999).

### DSF Determination of Ligand Binding Affinity

BSA binding affinity to octanoic acid (OA) was determined to confirm that DSF estimates for BSA binding were comparable to previously published values. Representative analysis of changes in BSA melting temperature induced by increasing concentrations of OA (0 to 5.5 mM) is shown in Figure 2A. The calculated K_d_ value (± SEM) of 3.13 ± 0.05 mM was in the expected range for fatty acid binding by bovine serum albumin (Figure 2B; Spector 1975).

**Figure 2.**
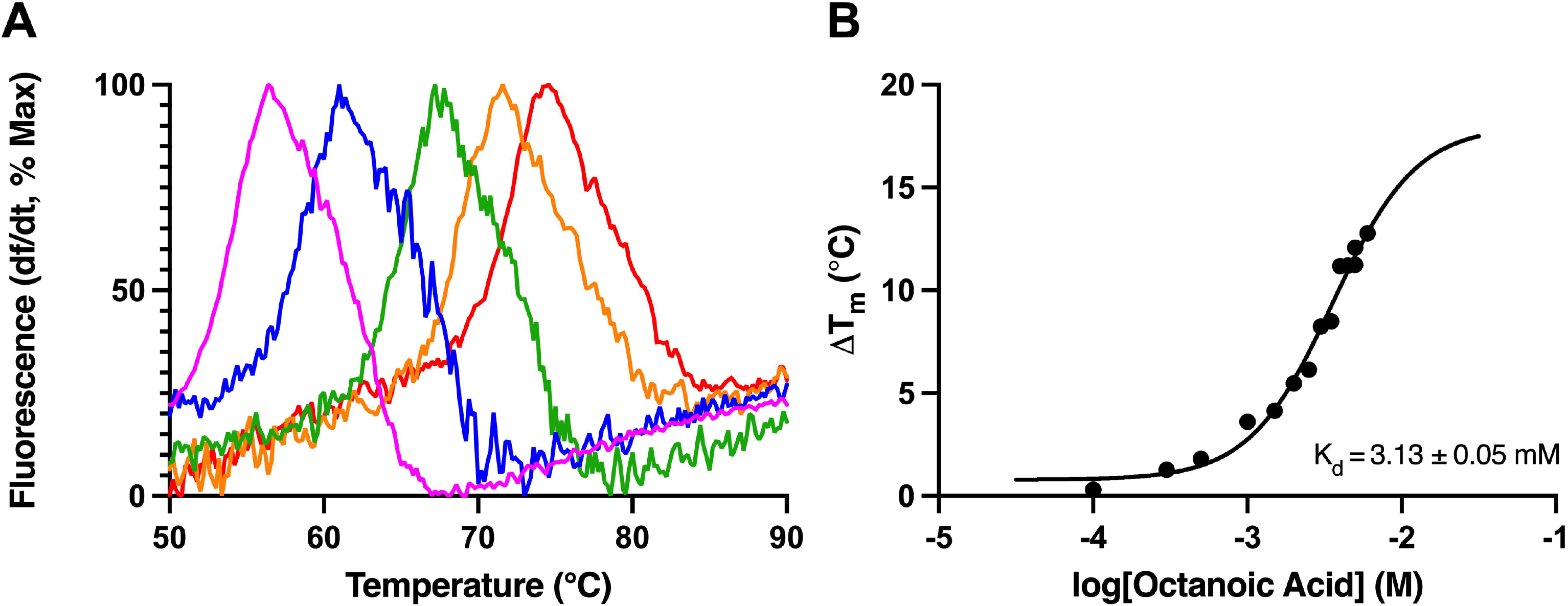
DSF analysis of BSA binding of octanoic acid. (A) Melt curves depicting changes in fluorescence for BSA with increasing concentrations of octanoic acid, normalized to percentage maximum fluorescence. Curves are plotted as the derivative of fluorescent signal divided by the derivative of time, as temperature is increased from 50 to 90°C. Increasing concentrations of octanoic acid, from 0 (pink) to 5.5 mM (red) are depicted. (B) The change in melting temperature (ΔT_M_) of BSA as a function of the log-transformed concentration of octanoic acid from 0 to 0.006 M. N = 6 replicates.

For each albumin, DSF was used to detect differences in T_M_ with increasing concentrations of each PFAS congener, and to estimate binding affinity values, reported in Table 2. Maximum ΔT_M_ values ranged from 3.53 to 15.9°C in HSA, 10.5 to 17.2°C in BSA, 5.74 to 13.9°C in PSA, and 4.76 to 12.2°C in RSA (Table 2). Binding affinities of PFAS ranged from 0.41 to 2.64 mM for HSA, 0.80 to 3.06 mM in BSA, 0.37 to 2.12 mM for PSA, and 0.33 to 1.90 mM for RSA (Table 2). Shown in Figure 3 are representative concentration-response analyses determining PFBS binding by HSA (Fig. 3A-B), BSA (Fig. 3C-D), PSA (Fig. 3E-F), and RSA (Fig. 3G-H). Each albumin bound PFAS congeners with charged functional groups, as evidenced by concentration dependent PFAS-induced shifts in protein melting temperature (Table 2). Consistent with previous findings, no changes in albumin T_M_ were detected for any concentration of 6:2 FTOH for any species (Jackson et al., 2021; Table 2). A unique biphasic melt curve in thermal stability change was exhibited for BSA binding of PFOS (Fig. 4A-H), a concentration response pattern consistent with unique binding interactions between PFOS and the BSA protein.

**Figure 3.**
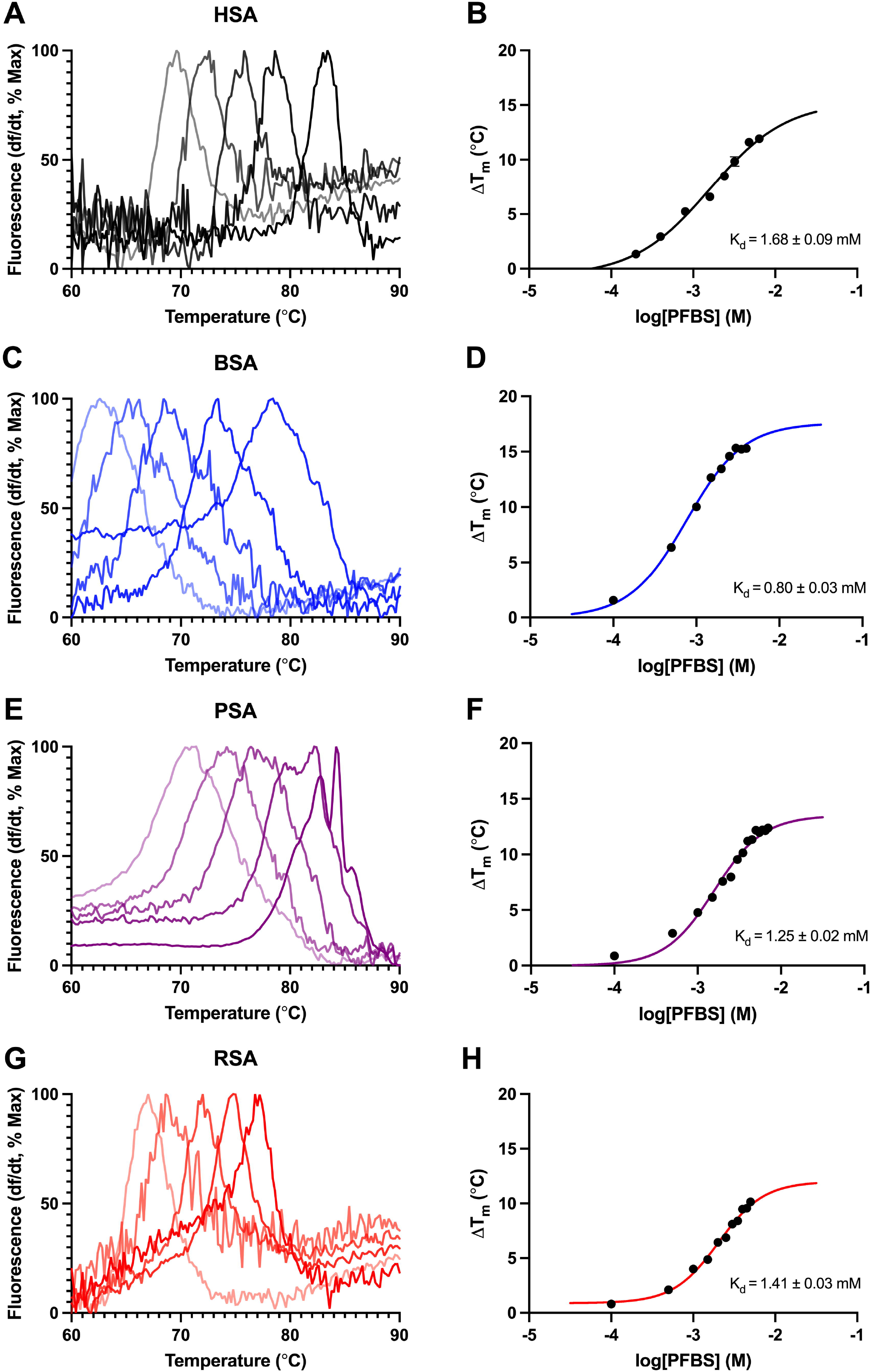
Melting temperature and concentration response analysis serum of albumin binding of PFBS. For each of the left-hand panels each curve depicts the change in fluorescent signal divided by change in time and is normalized to the percent maximum fluorescence for each albumin (A) HSA, (C) BSA, (E) PSA, and (G) RSA. Increased concentrations of PFBS are depicted by increased saturation of line color, as the temperature increased from 60 to 90°C. The change in melting temperature (ΔT_M_) as a function of the log-transformed concentration of PFBS (M) and calculated binding affinity (K_d_) is shown in panels (B) HSA, (D) BSA, (F) PSA, and (H) RSA. N = 6 replicates.

**Figure 4.**
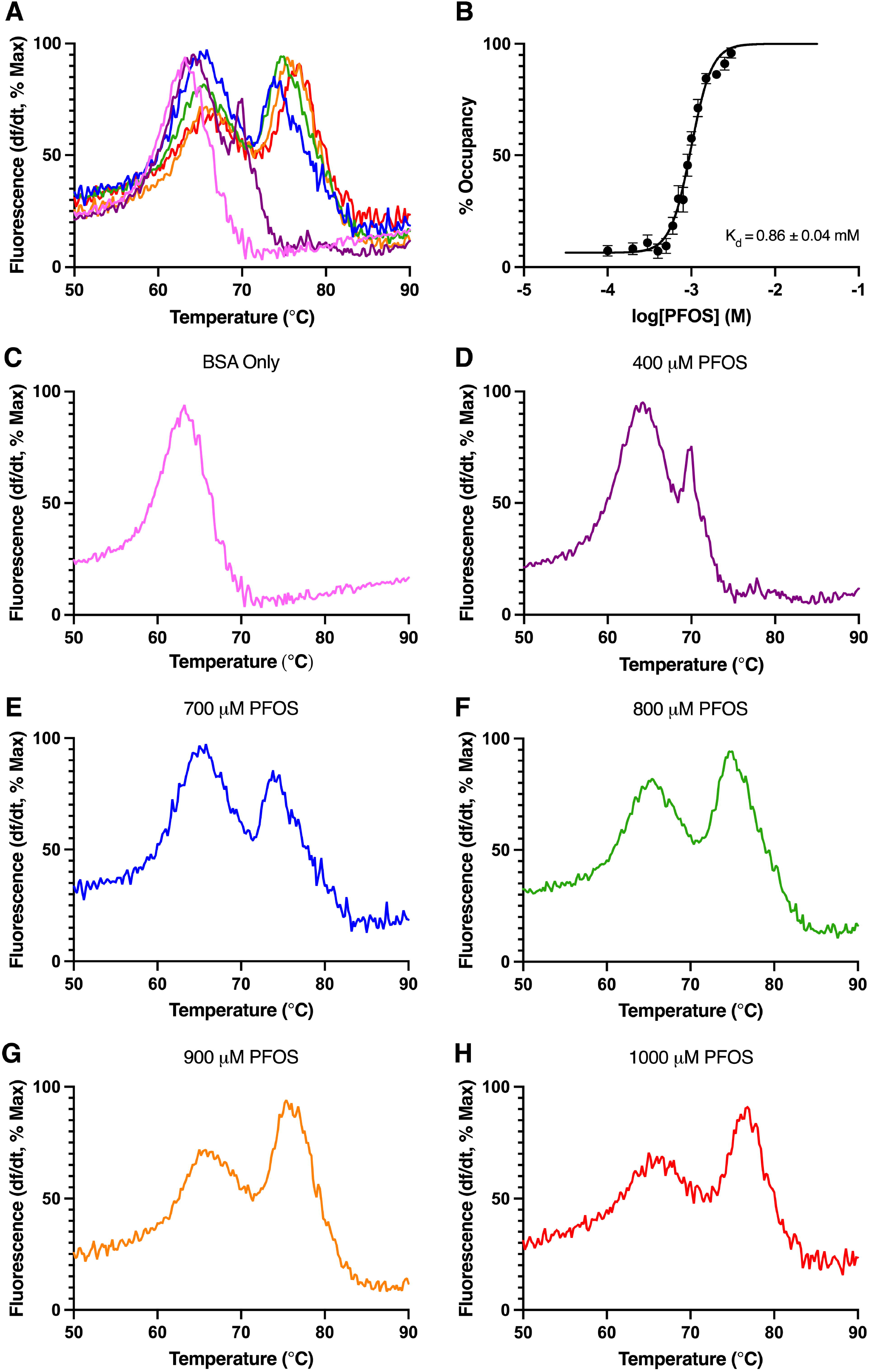
BSA binding of PFOS. (A) Change in fluorescence for BSA with increasing concentrations of PFOS. Increasing concentrations of PFOS are depicted by increasing wavelength of color from violet (0.4 mM) to red (1 mM). (B) The percent occupancy of BSA binding to PFOS, as a function of the log-transformed concentration of PFOS. (C-H) Melt curves for BSA at individual concentrations of PFOS from 0 to 1000 μM. N = 9 replicates.

**Table 2.** Comparison of PFAS binding affinities across species.

| Chemical | Chain Length | HSA |  |  | BSA |  |  |  | PSA |  |  |  | RSA |  |  |  |
| --- | --- | --- | --- | --- | --- | --- | --- | --- | --- | --- | --- | --- | --- | --- | --- | --- |
| | | $\Delta T_m$<br>(°C) | $K_d$<br>(mM) | $R^2$ | $\Delta T_m$<br>(°C) | $K_d$ (mM) | $R^2$ | d | $\Delta T_m$<br>(°C) | $K_d$ (mM) | $R^2$ | d | $\Delta T_m$<br>(°C) | $K_d$ (mM) | $R^2$ | d |
| PFBA | 4 | 6.43<br>(0.43) | 2.64<br>(0.22) | 0.96 | 10.5<br>(0.18) | 3.06<br>(0.27) | 0.93 | ns | 6.89<br>(0.07) | **1.28\***<br>**(0.03)** | 0.89 | **1.08** | 7.06<br>(0.09) | 1.90<br>(0.09) | 0.92 | ns |
| PFHxA | 6 | 10.5<br>(0.14) | 1.71<br>(0.22) | 0.98 | 17.1<br>(0.14) | **2.17\***<br>**(0.06)** | 0.98 | **0.88** | 13.9<br>(0.14) | **0.95\*\*\***<br>**(0.02)** | 0.97 | **2.16** | 12.2<br>(0.09) | 1.33<br>(0.02) | 0.97 | ns |
| PFOA | 8 | 13.0<br>(0.41) | 0.79<br>(0.09) | 0.97 | 14.4<br>(0.19) | **0.99\***<br>**(0.01)** | 0.92 | **2.12** | 9.68<br>(0.07) | **0.37\*\*\***<br>**(0.01)** | 0.89 | **7.12** | 8.0<br>(0.13) | **0.40\*\*\***<br>**(0.01)** | 0.97 | **0.89** |
| PFBS | 4 | 11.7<br>(0.21) | 1.68<br>(0.09) | 0.98 | 15.3<br>(0.12) | **0.80\*\*\***<br>**(0.03)** | 0.99 | **6.02** | 12.2<br>(0.06) | **1.25\*\*\***<br>**(0.02)** | 0.95 | **2.41** | 10.1<br>(0.08) | **1.41\*\***<br>**(0.03)** | 0.96 | **1.01** |
| PFOS | 8 | 15.9<br>(0.63) | 0.69<br>(0.02) | 0.93 | 15.3<br>(0.23) | 0.86<br>(0.05) | 0.79 | ns | 5.74<br>(0.20) | **0.38\***<br>**(0.06)** | 0.85 | **2.98** | 4.76<br>(0.38) | 0.42<br>(0.10) | 0.87 | ns |
| HFPO-DA | 5 | 13.7<br>(0.09) | 1.57<br>(0.03) | 0.95 | 12.7<br>(0.11) | **2.55\*\*\***<br>**(0.13)** | 0.95 | **1.99** | 10.0<br>(0.15) | **2.12\*\***<br>**(0.03)** | 0.93 | **1.23** | 7.56<br>(0.10) | 1.64<br>(0.05) | 0.94 | ns |
| 6:2 FTOH | 6 | N/A | N/A | N/A | N/A | N/A | N/A | N/A | N/A | N/A | N/A | N/A | N/A | N/A | N/A | N/A |
| 6:2 FTSA | 6 | 3.53<br>(0.26) | 0.41<br>(0.04) | 0.82 | 17.2<br>(0.19) | **1.39\*\*\***<br>**(0.04)** | 0.81 | **15.3** | 8.72<br>(0.09) | 0.52<br>(0.03) | 0.93 | ns | 7.99<br>(0.10) | 0.33<br>(0.02) | 0.99 | ns |

Maximum change in melting temperature values, represented as $\Delta T_m$ ( ± SEM) (°C), and protein binding affinities for each congener, represented as $K_d$ ( ± SEM) (mM). Bolded $K_d$ values represent values significantly different from HSA in GLM with Sidak's multiple comparisons test. $p < 0.05 = *$ , $p < 0.01 = **$ , $p < 0.001 = ***$ . N/A denotes non-binding. For comparisons in which significant differences were observed, Cohen's d (d) effect size values are reported, with bolded values denoting a large effect size. ns denotes a non-significant result. Each compound was run on at least 2 separate plates with $n \geq 5$ .

### Interspecies Comparison in Binding Affinities

Interspecies differences in calculated K_d_ values were evaluated with GLM, which revealed an overall effect of PFAS that accounted for 21.3% of model variance (χ^2^ = 815, p < .0001). Differences of species accounted for 5.72% of variance (χ^2^ = 197, p < .0001), and the interaction between PFAS and species accounted for 57.5% of variance (χ^2^ = 149, p < .0001).

Human albumin binding affinity for PFBA was decreased 69.4% compared to PSA (t = - 3.04, p = .016; Table 2), but no differences in binding affinity were detected for BSA (t = 1.22, p = .55) or RSA (t = -1.84, p = .21). Compared to HSA, binding affinity of PSA for PFHxA was increased 57.1% (t = -4.4, p = .00063) and BSA affinity was decreased 23.7% (t = 3.08, p = .016; Table 2), but there were no observed differences for RSA binding (t = -2.36, p = 0.079). Overall effects of protein were also detected for PFOA binding. Compared to HSA, PSA binding affinity for PFOA was increased 72.4% (t = -6.82, p < .0001), and RSA binding affinity was increased 65.5% (t = -5.49, p < .0001). By contrast, BSA binding affinity of PFOA was 22.5% lower than HSA (t = 3.21, p = .011).

Human albumin bound PFBS with 71.0% lower affinity (t = -10.6, p < .0001) compared to BSA, 29.4% lower affinity compared to PSA (t = -5.89, p < .0001), and 17.5% lower affinity compared to RSA (t = -3.68, p = .004; Table 2). Porcine albumin binding affinity of PFOS was increased 57.9% compared to HSA (t = -2.72, p = .032; Table 2), while no differences in PFOS binding affinity were detected for BSA (t = 1.41, p = .43) or RSA (t = -2.38, p = .072) compared to HSA.

Binding affinity of HSA for the perfluoroalkyl ether (PFAE) congener, HFPO-DA, was 47.6% greater than BSA (t = 6.86, p < .0001), and 29.8% greater than PSA (t = 3.79, p = .0028; Table 2), but was not different in RSA (t = .45, p = .96). The relative affinity of HSA for 6:2 FTSA, a fluorotelomer sulfonic acid, was increased 108.9% compared to BSA (t = 14.7, p < .0001; Table 2), whereas differences in 6:2 FTSA binding affinity were not detected for PSA (t = 2.31, p = .09) or RSA (t = -1.63, p = .31).

Comparing the impacts of chain length for a given subclass, GLM revealed significant differences in all species for albumin binding affinity of 4 and 8 carbon PFCA and PFSA congeners. No differences were detected for BSA binding of PFSAs (t = -.37, p = 1.0). The affinity of HSA binding of PFOA was 107.9% greater than PFBA (t = 7.0, p < .0001), and 83.5% greater for PFOS than PFBS (t = 4.01, p = 0.034; Table 3). For BSA (t = 8.36, p < .0001) and PSA (t = 4.07, p = .028), binding affinity for PFOA was 102.2% and 110.3% greater than PFBA, respectively (Table 3), with a similar 106.7% increase in PSA affinity for PFOS compared to PFBS (t = 3.97, p = .04). For RSA, affinity for PFOA was 130.4% greater than for PFBA (t = 4.69, p = .0021), and binding affinity for PFOS was 108.2% greater than PFBS (t = 4.50, p = .0049; Table 3).

**Table 3.**
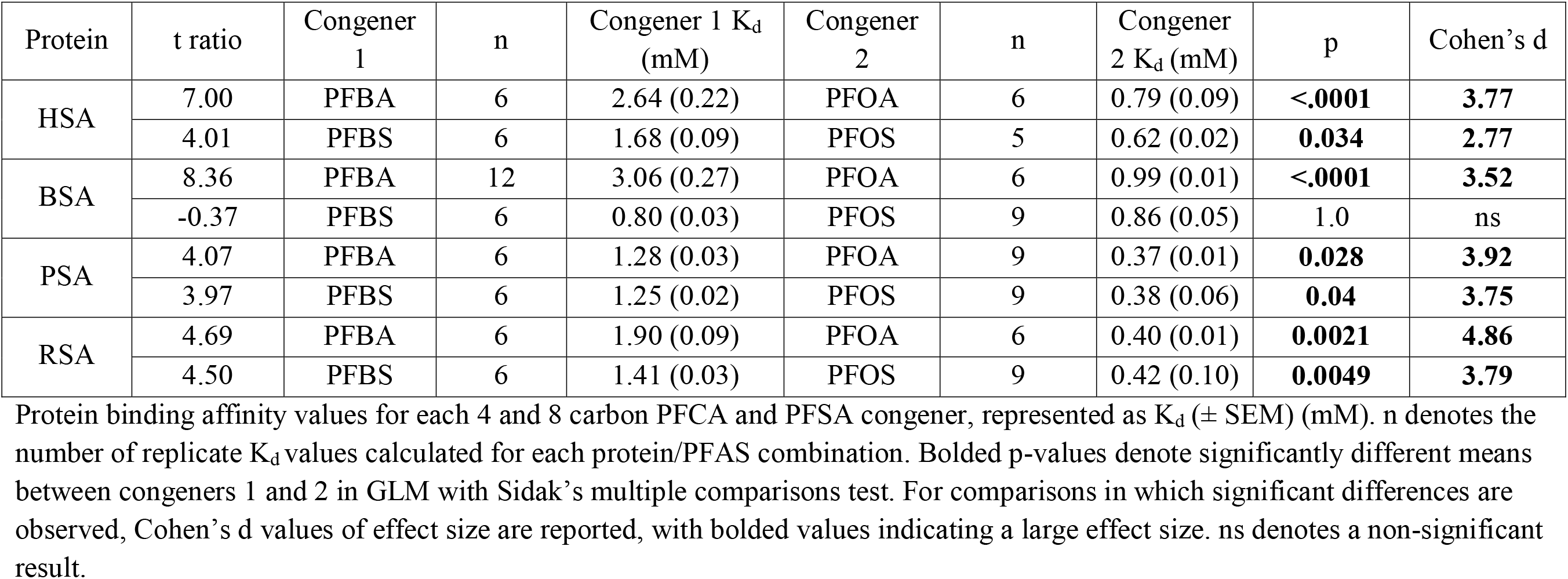
Comparison of 4 and 8 carbon PFAA binding affinities within species.

### Interspecies Amino Acid Sequence Alignment

SeqAPASS level 1 analysis, which compares primary sequence alignment, showed that the complete primary amino acid sequence of HSA was 79.5% similar to BSA, 78.5% similar to RSA, and 77.8% similar to PSA, with each albumin predicted to have similar ligand-binding interactions compared to HSA (Table 4, Figure S1).

**Table 4.** SeqAPASS level 1 and 3 analysis results.

| Albumin Species | % Similarity Overall | Sudlow Site I<br>Susceptibility Match | Sudlow Site II<br>Susceptibility Match | Trp214 Susceptibility<br>Match |
| --- | --- | --- | --- | --- |
| BSA | 79.5% | No | Yes | Yes |
| RSA | 78.5% | Yes | Yes | Yes |
| PSA | 77.8% | Yes | Yes | Yes |

Percentage of overall amino acid sequence similarity to HSA and evaluation of comparative susceptibility for chemical interactions at key binding sites. “Yes” denotes binding interactions predicted to be similar to HSA, “No” denotes binding interactions predicted to be dissimilar to HSA.

Key amino acid residues involved in polar and electrostatic interactions between HSA and PFAS were further evaluated using level 3 analysis, which compares individually selected amino acid residues. Level 3 analysis was constrained to include residues in two well-characterized binding sites, Sudlow sites I and II, which are involved in HSA binding of drugs, fatty acids, and some PFAS (Sudlow et al. 1975; Sudlow et al. 1976; Chen and Guo 2009; Ascenzi and Fasano 2010). The amino acid residues of the Sudlow II sites of HSA, PSA, BSA, and RSA were identical, demonstrating strong structural and functional conservation across species (Table 4, S1, S2). Sudlow site I was more divergent relative to HSA; however, PSA and RSA were predicted to have chemical binding capabilities similar to HSA due to the functional conservation of amino acid substitutions (Table S1). In contrast, Sudlow site I of BSA, which contains four functionally conserved amino acid substitutions and a F211L substitution expected to alter functionality of the protein, was predicted to have different binding capabilities than HSA. Computational substitution of the phenylalanine residue in the Sudlow I site of HSA with a leucine residue (F211L), representing a key amino acid substitution found in BSA, was predicted to be destabilizing to HSA with a -1.97 kcal/mol enthalpy change (ΔΔG).

### Molecular Docking Predictions

To assess concordance of experimentally derived albumin-PFAS binding affinity analysis and computationally predicted values, Autodock Vina was used to predict enthalpy changes in the free energy of binding (ΔG, kcal/mol) for three different HSA structural conformations and one BSA structure for 7 PFAS congeners analyzed (Table S3). Pearson’s correlation was calculated to assess the linear relationship between experimental K_d_ values and molecular docking ΔG predictions for each PFAS-protein combination (Fig. 5). Across HSA structures, strong positive correlations between variables were found for conformations 1E7G (Fig. 5A; r(12) = .93, p = .0021), 4E99 (Fig. 5B; r(12) = .92, p = .0031), and 1H9Z (Fig. 5C; r(12) = .91, p = .0039). With the exception of PFBS, there was also a strong correlation between the variables for all tested congeners in 4F5S, the BSA conformer, (Fig. 5D; r(10) = .98, p = .0006).

**Figure 5.**
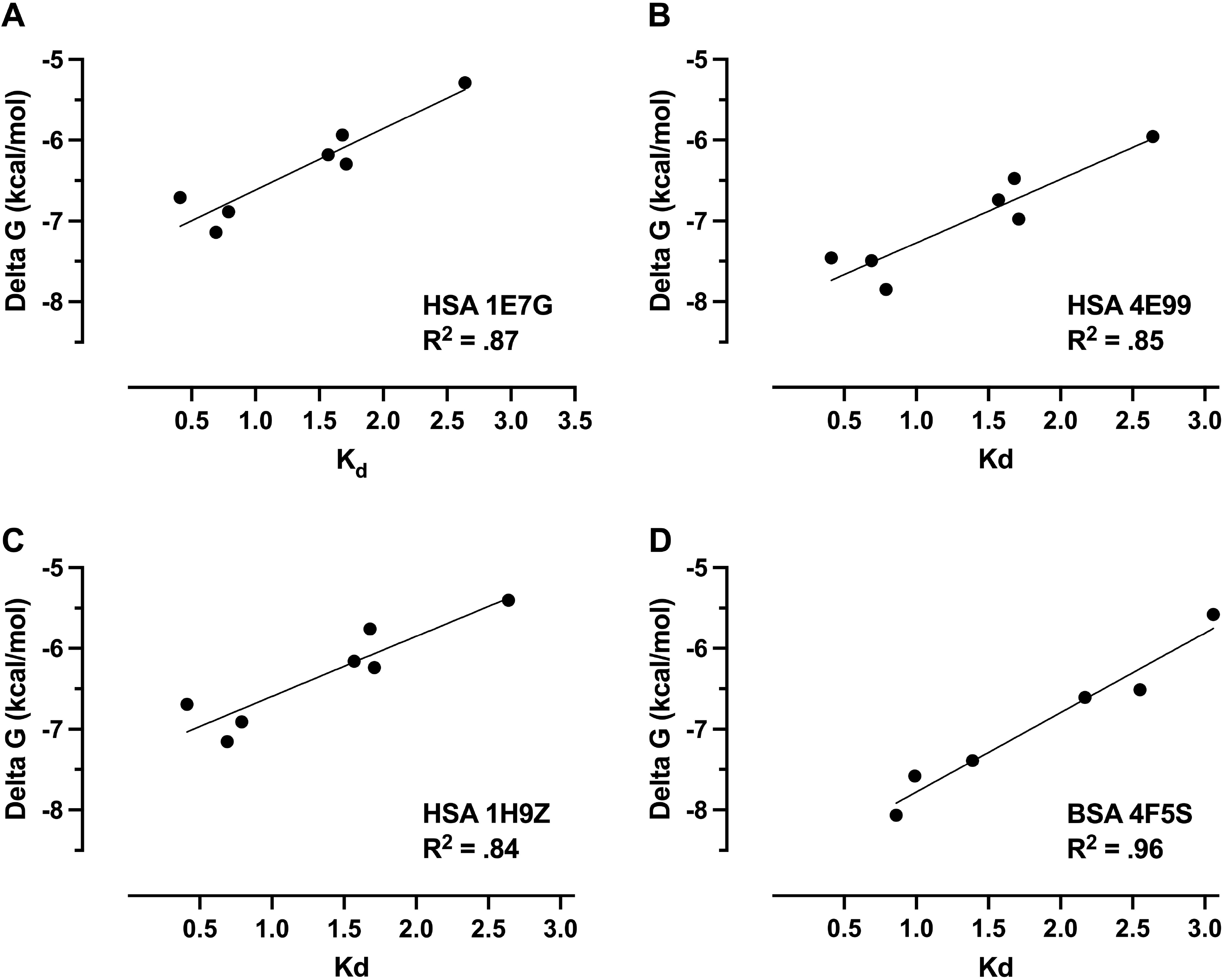
Correlations between DSF-measured dissociation constants (K_d_) and computational predictions of ΔG of binding different PFAS congeners. (A) HSA 1E7G (conformation bound to myristic acid), (B) HSA 4E99 (conformation bound to PFOS), (C) HSA 1H9Z (intermediate conformation), and (D) BSA 4F5S (native protein with no ligand) without PFBS.

## Discussion

A DSF assay was used in this study to compare interspecies differences in albumin binding for an environmentally relevant set of PFAS congeners, to fill critical data gaps and decrease uncertainty in animal to human extrapolation. Protein binding affinities for PFAS are an important determinant of bioaccumulative potential, and a lack of understanding of serum albumin interactions with a majority of PFAS congeners is a major data gap in the field (Hall and Peng 2020; Cheng et al. 2021). Most experimental research used to understand PFAS accumulation and toxicity has been conducted in animal models, presenting a need for extrapolation of data to predict human outcomes. Important improvements have been made in the accuracy of computational predictions of toxicity-related factors, however absence of experimental data for many chemicals remains a challenge to the development of *in silico* frameworks that improve translational predictive capability (Cheng and Ng 2017; Chou and Lin 2019; Cheng et al. 2021; Chou and Lin 2021; Deepika et al. 2021). Specifically, the lack of fundamental physiochemical data remains a major limitation in model performance, particularly for the 1000s of PFAS congeners lacking experimental data.

Our analyses demonstrated that each PFAS congener that was bound by human serum albumin was also bound by bovine, porcine and rat albumin. Those results agreed with sequence homology predictions indicating that HSA, BSA, PSA, and RSA would have similar ligand-binding properties. Although data reporting affinities of BSA and RSA for PFAS are limited, the K_d_ estimates derived using our high-throughput DSF platform were within the same order of magnitude as published albumin-PFAS binding affinities (Klevens and Ellenbogen 1954; Han et al. 2003; MacManus-Spencer et al. 2010; Jackson et al. 2021). It is notable that absolute binding affinities vary greatly across studies due to differences in experimental conditions. As expected, our DSF-determined binding affinities were lower than those reported in studies using other methods; those findings are due to the fact that DSF estimates binding affinities over a range of temperatures, whereas most other approaches estimate binding affinity at a single, typically lower temperature (Vivoli et al. 2014; Jackson et al. 2021).

Confirming our findings that the fluorotelomer alcohols were not bound by HSA, the three additional mammalian serum albumins likewise did not bind 6:2 FTOH. We interpret this lack of observed binding to support the essential influence of a charged functional headgroup as a key structural determinant of albumin-PFAS binding (Jackson et al. 2021). Multiple lines of evidence support this conclusion, including *in vivo* studies that found 8:2 FTOH did not accumulate in pig or rat tissues after oral exposure (Fasano et al. 2006; Xie et al. 2020).

Protein binding is an important mediator of PFAS transport to blood-rich tissues, where equilibrium gradients can drive tissue uptake and resulting toxicity (Ng and Hungerbühler 2014; Ng and Hungerbühler 2015; Cheng and Ng 2017). Fluorotelomer alcohol congeners and other PFAS that are poorly bound by serum proteins are expected to have important differences in toxicokinetic behaviors, including more rapid metabolism to breakdown products, shorter biological half-lives, differences in accumulation in body compartments, and changes in observed elimination rate. Such impacts are demonstrated by the absence of 8:2 FTOH accumulation in pig and rat tissues (Fasano et al. 2006; Xie et al. 2020). Further, Xie et al. (2020) found that while 8:2 FTOH did not bioaccumulate, metabolic breakdown products of 8:2 FTOH with charged carboxylic headgroups accumulated in tissues of exposed pigs. Our results demonstrating that HSA, BSA, PSA, and RSA bind to 6:2 FTSA but not 6:2 FTOH, along with *in vivo* evidence that 8:2 FTOH does not accumulate in rat and pig tissues, provides strong evidence that PFAS containing a charged functional group have increased bioaccumulative potential in mammalian species compared to their uncharged analogues (Fasano et al. 2006; Xie et al. 2020).

In addition to functional groups, perfluorinated carbon chain length is a key physiochemical property that determines relative HSA binding affinity for PFAS (Bischel et al. 2010; Ng and Hungerbuehler 2015; Liu et al. 2017; Jackson et al. 2021; Crisalli et al. 2023). Numerous studies have found that HSA affinities for linear PFAAs of increasing chain length display a U-shaped trend with optimal binding between 6-9 aliphatic carbons, while congeners with fewer (4-5) or more (10-12) aliphatic carbons are bound less tightly (Salvalaglio et al. 2010; Ng and Hungerbühler 2014; Jackson et al. 2021). To investigate the hypothesis that BSA, PSA, and RSA would also bind optimally to PFAAs with 6-9 aliphatic carbons, we compared relative affinities for PFCA and PFSA congeners with 4 and 8 aliphatic carbons. Binding affinity was increased for PFOA compared to PFBA in all tested albumins, and for all species except BSA, PFOS was bound with greater affinity than PFBS. Those results reveal interspecies conservation of the relationship between carbon chain length and albumin binding affinity, and highlight perfluorinated carbon chain length as a critical physiochemical determinant of albumin binding.

Comparative interspecies albumin-PFAS binding data reported in this study offer insight into the suitability of uncertainty factors (UFs) in the estimation of human health exposure limits. In risk assessment, UFs are used to account for the uncertainty of interspecies variability when extrapolating findings from experimental animal research for human relevance, and require an understanding of PFAS toxicokinetic differences across species to best protect human health (Dankovic et al. 2015). However, in the absence of species or chemical-specific data, an uncertainty factor of 10 is typically used to extrapolate from animal to human data (Dourson et al. 1996; Dankovic et al. 2015). Dissimilar to all other congeners tested, 6:2 FTOH was not bound by albumin of any species, demonstrating that the use of a single uncertainty factor for all PFAS is not appropriate and that chemical or class-specific and data-derived values may be necessary for some pharmacokinetic parameters.

Beyond the lack of FTOH binding, more subtle differences in species-specific binding affinity emerged. Comparing albumin binding patterns for PFAAs to HSA, our data revealed that PSA and RSA bound these subclasses with increased affinities, whereas BSA bound with decreased affinities, with the notable exception of PFBS (Fig. 6). In contrast, relative binding affinity for the perfluoroalkyl ether (HFPO-DA) was greatest for HSA, with decreased affinities observed for PSA and BSA binding (Fig. 6). Differences in protein flexibility likely explain some binding affinity differences, as albumin undergoes conformational changes to accommodate ligand binding. For example, decreased affinities seen in BSA may be partly explained by the increased rigidity of BSA compared to HSA, which confers less flexibility to accommodate binding (Ketrat et al. 2020).

**Figure 6.**
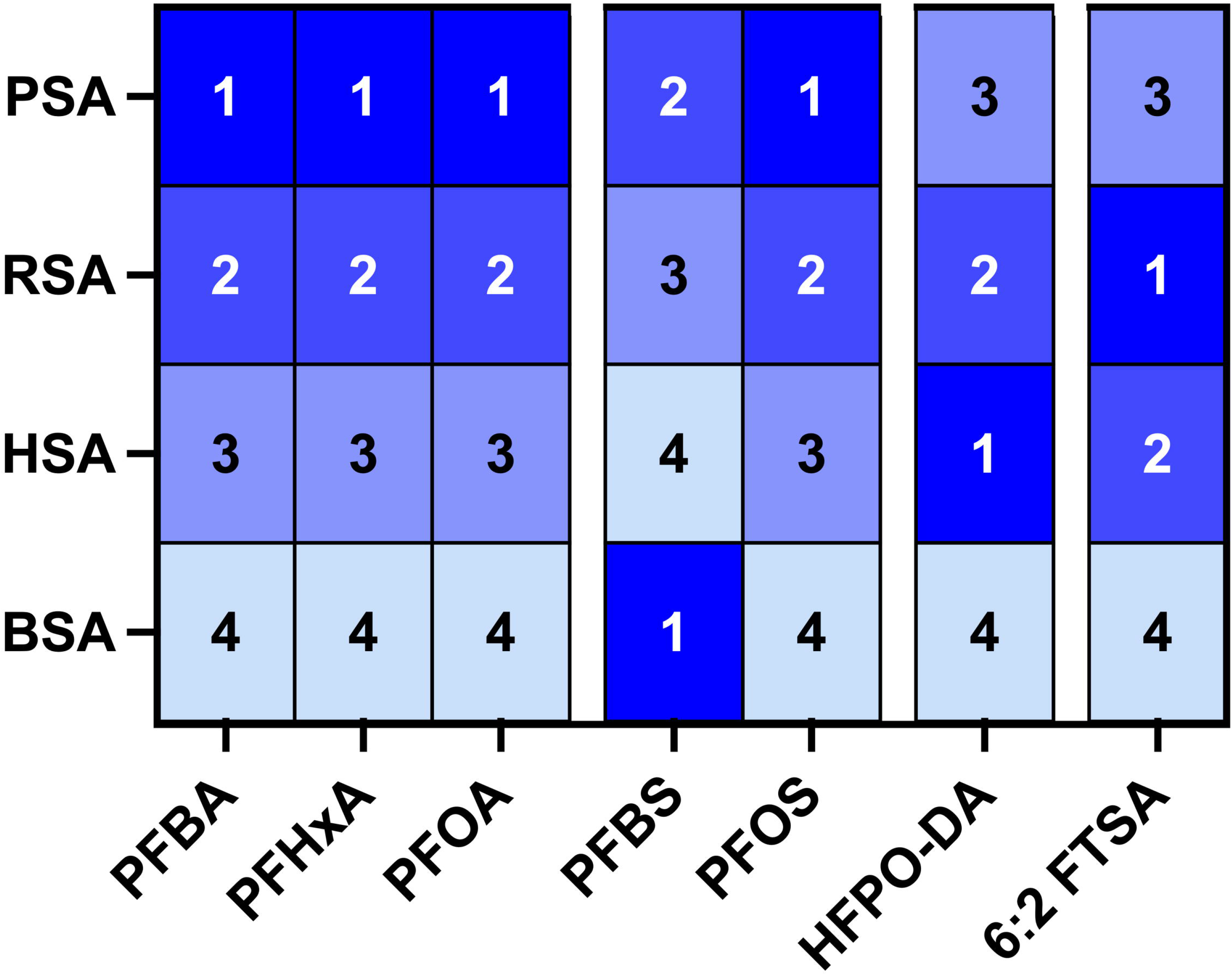
Heatmap showing rank order of binding affinities for each bound PFAS congener. For each congener 1 indicates highest relative affinity binding and 4 the lowest affinity for each species.

A notable difference in thermal melt profiles across species was found in BSA-PFOS binding where a biphasic denaturation profile was observed, a profile not seen for other albumin proteins or for BSA with other analyzed PFAS congeners. BSA melted at two different temperatures in the presence of PFOS, demonstrating saturation of binding sites (Vivoli et al. 2014). With increasing PFOS concentrations, the proportion of BSA melting at the baseline ligand-free melting temperature decreased, and the remaining proportion of BSA melting at an elevated temperature increased. Biphasic denaturation profiles were also observed in differential scanning calorimetry experiments measuring BSA binding of surfactants (sodium dodecyl sulfate and a dirhamnolipid biosurfactant) (Giancola et al. 1997; Deep and Ahluwalia 2001; Sánchez et al. 2008), and for HSA binding of palmitic acid (Brandt and Andersson 1976; Shrake and Ross 1988; Nemergut et al. 2023). While several hypotheses have been proposed, there is no clear consensus as to the cause of biphasic melt curves across protein-ligand affinity studies (Shrake and Ross 1988; Giancola et al. 1997; Luan et al. 2014; Nemergut et al. 2023). Due to the heterogeneity of binding sites across the three homologous domains of the albumins studied, with each site comprising different chemical properties that are optimally suited for different ligands, we hypothesize that PFOS is preferentially binding and stabilizing one or two of the homologous domains of BSA (Fasano et al. 2005; Nemergut et al. 2023).

Observed binding affinity differences are also likely dictated by changes in the amino acid sequence of each albumin, particularly in binding sites. However, details regarding site-specific albumin binding affinity for different PFAS congeners are poorly understood. Albumin binds to several PFAA congeners through non-covalent hydrophobic and electrostatic interactions (Fasano et al. 2005; Zhang et al. 2009; Salvalaglio et al. 2010; Liu et al. 2017; Chen et al. 2020). Proposed albumin binding sites for PFAS include Sudlow sites I and II, interactions impacting Trp214, and several fatty acid binding sites (Chen and Guo 2009; D’eon et al. 2010; MacManus-Spencer et al. 2010; Salvalaglio et al. 2010; Sheng et al. 2020; Maso et al. 2021; Cao et al. 2022; Moro et al. 2022). Analysis of binding affinity data based on structural changes of HSA Trp214 residue via quenching of intrinsic tryptophan fluorescence, and Sudlow sites I and II via fluorescence displacement of site-specific probes, found that carbon chain length and functional group of PFAA congeners strongly influenced HSA binding capability at each site (Chen and Guo 2009). While our DSF assay does not have site-specific experimental resolution, our differential binding, homology, and docking results support the fact that subtle changes in amino acid sequence in putative PFAS binding sites impact albumin binding affinities for PFAS.

Molecular docking was utilized to further investigate HSA and BSA binding of PFAS, and to compare the dissociation constants determined from our *in vitro* assay results with *in silico* binding predictions from Autodock Vina, which was used to simulate albumin-PFAS binding at 26.85°C. Like results reported by Ng and Hungerbühler (2015), predicted binding affinities were increased compared to those determined experimentally. The use of multiple HSA structures allowed binding affinity assessment across different conformations, addressing the limitation that Autodock Vina does not account for changes in protein conformation during ligand binding (Trott and Olson 2009; Ng and Hungerbuehler 2015). Across HSA structural conformations, there was a strong positive correlation between experimentally determined dissociation constants and docking predictions. Except for PFBS, there was also a strong positive correlation between these values for BSA binding. Strong agreement of experimental and predicted values provides two lines of evidence to support our experimental conclusions and strengthens the hypothesis that subtle interspecies differences in PFAS interactions with amino acids in specific binding sites will influence PFAS binding affinities.

There was poor concordance between the molecular docking prediction for BSA-PFBS binding compared to our experimentally determined affinity, with molecular docking predicting a much lower binding affinity. Tight binding of BSA to PFBS, comparable to binding affinity for PFOS, was also reported in a study utilizing equilibrium dialysis to determine protein-water partition coefficients (Bischel et al. 2011). As seen in our experiments, Bischel et al. (2011) also found significantly increased BSA binding affinities for PFCAs with increased aliphatic chain lengths, but not PFSAs. In contrast, two other studies on BSA-PFAS binding reported decreased affinity for PFBS compared to PFOS (Allendorf et al. 2019; Alesio et al. 2022). However, Bischel et al. (2011) states that some common fluorescence methods may not be suitable for assessing albumin binding to short chain PFAAs, including the fluorescence quenching approach used by Alesio et al. (2022), due to the failure of these short-chain PFAAs to cause the conformational changes in secondary structure of BSA that elicit spectral changes (Hebert and MacManus-Spencer 2010; Qin et al. 2010). Inconsistency between our *in vitro* and *in silico* results for BSA-PFBS binding, as well as the biphasic denaturation profile seen only with BSA-PFOS binding, suggests that PFAS binding by BSA may involve major differences in binding modalities for sulfonic acids. These and other HSA-PFAS binding studies reporting different trends in binding affinities across different structural classes of PFAS (Jackson et al. 2021), or failure of proposed models of HSA-PFAA binding to generate accurate results for short-chain compounds (Hebert and MacManus-Spencer 2010), also concluded that differences in the mode of binding for these compounds are a likely explanation for divergence from predicted relative binding affinities.

At Sudlow site I of BSA, a single amino acid residue substitution was identified that was predicted by SeqAPASS to alter functionality of this key PFAS-binding site relative to HSA. Sudlow site I is found in albumin’s subdomain IIA, overlaps with fatty acid site 7, and serves as the warfarin binding site, in addition to binding other bulky, heterocyclic anionic compounds (Fasano et al. 2005; Ascenzi and Fasano 2010). In the mature HSA protein, the key residues involved in ligand binding evaluated at Sudlow site I included Tyr150, Glu153, Lys195, Gln196, Lys 199, Phe211, Trp214, Ala215, Arg218, Arg222, Leu238, His242, Arg257, His288, and Ala291 (Ascenzi and Fasano 2010). All of these key residues are functionally conserved in PSA and RSA with respect to HSA. However, the phenylalanine residue (Phe211) found in HSA is replaced with a leucine residue (Leu210) in BSA. Phenylalanine and leucine are both non-polar and hydrophobic, however phenylalanine is a bulkier, aromatic residue and is likely to introduce more steric hindrance at the binding site than leucine. Calculations using the DUET web-tool supported the SeqAPASS prediction of altered functionality at Sudlow site I of BSA, predicting that a F211L mutation in HSA would yield a destabilizing enthalpy change which would likely alter protein-ligand interactions at this site. Our findings suggest that this destabilizing substitution at Sudlow site I in BSA alters binding for ligands known to bind this site, including PFBA, PFBS, PFOA, and PFOS (Chen and Guo 2009).

Binding affinities of PSA for PFCAs and PFSAs were significantly greater than HSA affinities, while affinity for HFPO-DA was significantly decreased. Relative to HSA, PSA contains the greatest percentage of amino acid sequence divergence (77.8% homology). This degree of sequence divergence suggests that PSA should exhibit more functional differences compared to HSA than BSA or RSA, which agrees with our experimental findings; PSA binding affinities showed differences from HSA affinities with large effect sizes for all of the PFAA and PFAE congeners (Table 2).

To more fully understand interspecies differences in PFAS distribution and retention, it is necessary to consider other parameters in conjunction with albumin binding affinity, including binding affinities of other proteins, tissue transfer efficiencies, and metabolism (LaLone et al. 2013; Cheng and Ng 2017; Chou and Lin 2019; Gao et al. 2019). For many protein-bound ligands, including pharmaceuticals, increased protein binding affinity increases biological half-life (Sleep et al. 2013; Bech et al. 2018; Zorzi et al. 2019). It is more challenging to predict the impacts of albumin binding affinity on perfluorinated compounds because they are not metabolized, unlike most pharmaceuticals. Coupling our data with published findings, we hypothesized that increased binding affinity of PSA for PFAAs will increase PFAA half-life in the blood compartment of pigs. In support of this conclusion, a study conducted by Numata et al. (2014) found that blood plasma in pigs has higher affinity for PFAAs, relative to reported affinities for cows and sheep (Kowalczyk et al. 2013; van Asselt et al. 2013; Numata et al. 2014). However, PFBS is the only congener which has a longer elimination half-life in pigs compared to reported values in humans, while half-lives for PFOS, PFOA, and PFHxA are longer in humans, highlighting the need to consider more physiochemical parameters in addition to albumin binding (Table 5).

**Table 5.**
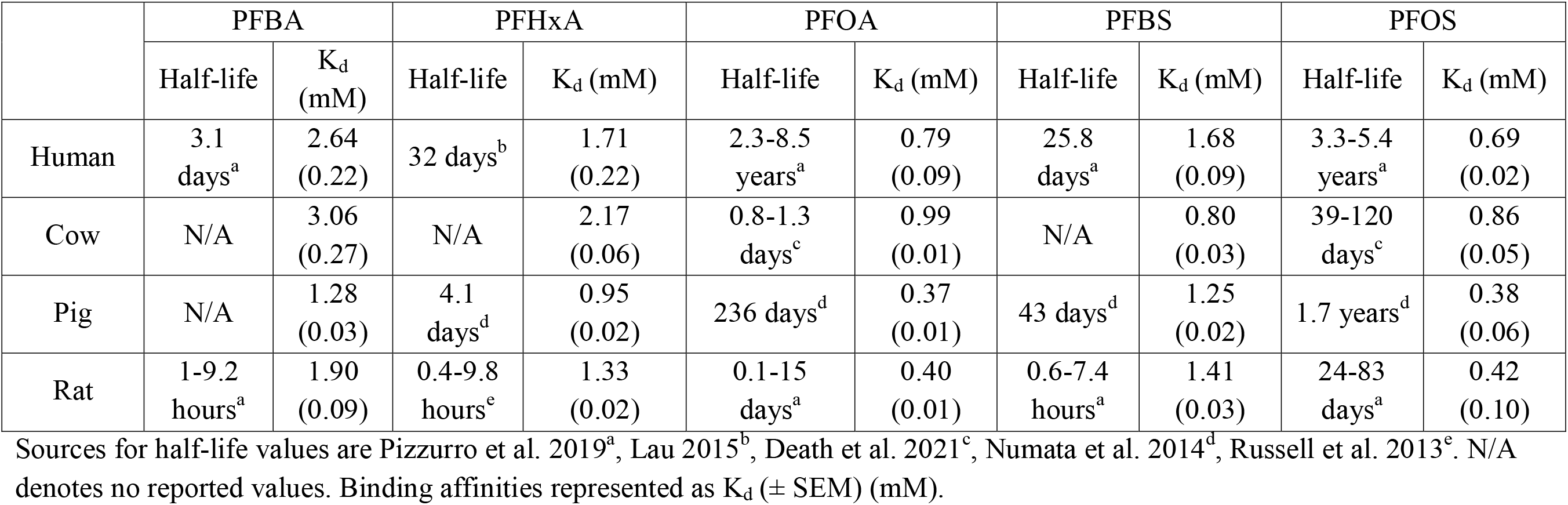
Reported elimination half-lives of PFAAs in serum/plasma across species and DSF-determined albumin binding affinities.

Understanding differences in PFAS-protein interactions between rats and humans is an important endpoint necessary for accurate translational extrapolation of experimental rodent toxicity studies. The observed differences in PFAS binding affinities between HSA and RSA in this study could serve as important data to decrease uncertainty in the application of animal model data for human relevance, and can be used to help understand differences in PFAS behavior across species. Overall, HSA and RSA were quite similar in binding affinities for most PFAS, and our findings suggest that guideline toxicity studies in rat models are not likely to be greatly impacted by species differences in albumin binding for the PFAS congeners in our test set. PFAS have also been detected widely in the tissues of livestock species, including cows and pigs, and are excreted in dairy milk (Guruge et al. 2008; van Asselt et al. 2013; Numata et al. 2014; Death et al. 2021; Rock et al. 2023). As one of the major routes of exposure to PFAS for humans is consumption of contaminated food and water, assessment of PFAS accumulation in livestock species is also needed to evaluate the role of livestock consumption in human PFAS exposure (Sunderland et al. 2019; Roth et al. 2020). Our findings demonstrate that HSA-PFAS binding is less similar to BSA and PSA, suggesting increased uncertainty in extrapolation of toxicokinetic parameters and distribution of PFAS in cows and pigs. Data generated in our study using the DSF assay provide necessary interspecies comparisons and serve as an essential step toward understanding the physiochemical properties of PFAS across species.

## Acknowledgments

We would like to sincerely thank Dr. Nicholas Harmer for troubleshooting assistance in our modeling of the biphasic melt curves, and Yuexin “Cara” Cao and Dr. Carla Ng for their expertise and guidance with molecular docking experiments.

