## Supplemental Tables S1-S3; Fig. S1 for "Quantitative Cross-Species Comparison of Serum Albumin Binding of Per- and Polyfluoroalkyl Substances from Five Structural Classes"

**Index:**

1. Table S1: SeqAPASS level 3 analysis for Sudlow Site I
2. Table S2: SeqAPASS level 3 analysis for Sudlow Site II
3. Table S3: Delta G Binding Predictions from Autodock Vina
4. Figure S1: Full protein sequence alignment

Supplementary Material

Table S1. SeqAPASS level 3 analysis for Sudlow Site I.

| Protein | NCBI Accession Number | Similar Susceptibility | Amino Acid 1 | Amino Acid 2 | Amino Acid 3 | Amino Acid 4 | Amino Acid 5 | Amino Acid 6 | Amino Acid 7 | Amino Acid 8 | Amino Acid 9 | Amino Acid 10 | Amino Acid 11 | Amino Acid 12 | Amino Acid 13 | Amino Acid 14 | Amino Acid 15 |
|---|---|---|---|---|---|---|---|---|---|---|---|---|---|---|---|---|---|
| HSA | AAA98797.1 | Yes | 150Y | 153E | 195K | 196Q | 199K | 211F | 214W | 215A | 218R | 222R | 238L | 242H | 257R | 288H | 291A |
| BSA | P02769.4 | No | 149Y | 152E | 194R | 195Q | 198R | 210L | 213W | 214S | 217R | 221K | 237L | 241H | 256R | 287H | 290A |
| RSA | NP_599153.2 | Yes | 150Y | 153E | 195R | 196Q | 199K | 211F | 214W | 215A | 218R | 222R | 238L | 242N | 257R | 288Q | 291A |
| PSA | ABM92961.1 | Yes | 149Y | 152E | 194K | 195Q | 198K | 210F | 213W | 214S | 217R | 221R | 237L | 241H | 256R | 287H | 291A |

Sequence alignment of amino acids in Sudlow site I, and SeqAPASS classification of susceptibility for ligand binding compared to HSA. Green boxes indicate identical amino acids compared to HSA, yellow boxes indicate partial (functionally conserved) matches to HSA, and red boxes indicate amino acids that do not match HSA.

Table S2. SeqAPASS level 3 analysis for Sudlow Site II.

| Protein | NCBI Accession Number | Similar Susceptibility | Amino Acid 1 | Amino Acid 2 | Amino Acid 3 | Amino Acid 4 | Amino Acid 5 | Amino Acid 6 | Amino Acid 7 |
|---|---|---|---|---|---|---|---|---|---|
| HSA | AAA98797.1 | Yes | 387L | 410R | 411Y | 414K | 433V | 453L | 489S |
| BSA | P02769.4 | Yes | 386L | 409R | 410Y | 413K | 432V | 452L | 488S |
| RSA | NP_599153.2 | Yes | 387L | 410R | 411Y | 414K | 433V | 453L | 489S |
| PSA | ABM92961.1 | Yes | 386L | 409R | 410Y | 413K | 432V | 452L | 488S |

Sequence alignment of amino acids in Sudlow site II, and SeqAPASS classification of susceptibility for ligand binding compared to HSA. Green boxes indicate identical amino acids compared to HSA.

Table S3. Delta G of Binding Predictions from Autodock Vina

| Ligand | Protein | PDB ID | Geometric Mean of ΔG (kcal/mol) | Geometric SD of ΔG (kcal/mol) |
|---|---|---|---|---|
| PFBA | HSA | 1E7G | -5.29 | 0.46 |
| PFBA | HSA | 4E99 | -5.95 | 0.26 |
| PFBA | HSA | 1H9Z | -5.40 | 0.77 |
| PFBA | BSA | 4F5S | -5.58 | 0.85 |
| PFHxA | HSA | 1E7G | -6.29 | 0.70 |
| PFHxA | HSA | 4E99 | -6.97 | 0.65 |
| PFHxA | HSA | 1H9Z | -6.24 | 1.03 |
| PFHxA | BSA | 4F5S | -6.61 | 1.40 |
| PFOA | HSA | 1E7G | -6.88 | 0.92 |
| PFOA | HSA | 4E99 | -7.85 | 0.92 |
| PFOA | HSA | 1H9Z | -6.91 | 1.02 |
| PFOA | BSA | 4F5S | -7.58 | 1.78 |
| PFBS | HSA | 1E7G | -5.94 | 0.68 |
| PFBS | HSA | 4E99 | -6.48 | 0.58 |
| PFBS | HSA | 1H9Z | -5.76 | 0.97 |
| PFBS | BSA | 4F5S | -6.10 | 1.39 |
| PFOS | HSA | 1E7G | -7.14 | 0.99 |
| PFOS | HSA | 4E99 | -7.49 | 1.11 |
| PFOS | HSA | 1H9Z | -7.15 | 1.36 |
| PFOS | BSA | 4F5S | -8.06 | 1.87 |
| 6:2 FTSA | HSA | 1E7G | -6.71 | 0.74 |
| 6:2 FTSA | HSA | 4E99 | -7.46 | 0.75 |
| 6:2 FTSA | HSA | 1H9Z | -6.69 | 0.96 |
| 6:2 FTSA | BSA | 4F5S | -7.39 | 1.72 |
| GenX | HSA | 1E7G | -6.18 | 0.94 |
| GenX | HSA | 4E99 | -6.74 | 0.65 |
| GenX | HSA | 1H9Z | -6.16 | 0.95 |
| GenX | BSA | 4F5S | -6.51 | 1.26 |

Geometric mean and standard deviation of Autodock Vina predicted ΔG of binding across six binding sites, in three high-resolution structural conformations of HSA and one of BSA.

Supplementary Material

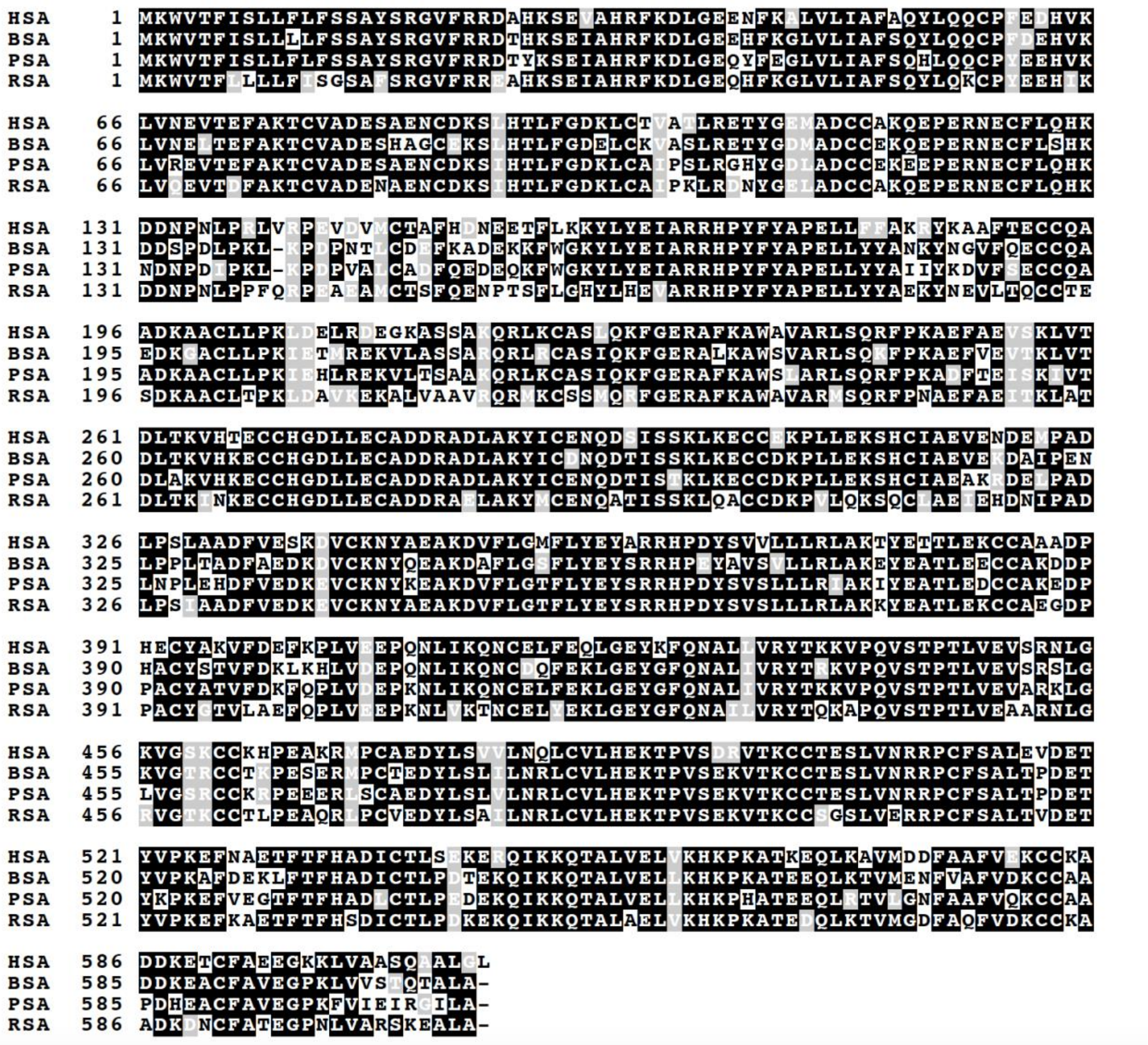


Figure S1. Full protein sequence alignment of the 609 amino acids in serum albumin, including the 24 amino acids in the prepro protein and the 585 amino acids in the mature albumin protein. Black background indicates identical residues, gray background indicates similar residues, and white background is used for remaining residues. Sequence alignment generated by T-Coffee (https://tcoffee.crg.eu/apps/tcoffee/do:regular).
